# A high-content screen reveals new regulators of nuclear membrane stability

**DOI:** 10.1101/2023.05.30.542944

**Authors:** Amanda L. Gunn, Artem I. Yashchenko, Julien Dubrulle, Jodiene Johnson, Emily M. Hatch

## Abstract

Nuclear membrane rupture is a physiological response to multiple *in vivo* processes, such as cell migration, that can cause extensive genome instability and upregulate invasive and inflammatory pathways. However, the underlying molecular mechanisms of rupture are unclear and few regulators have been identified. In this study, we developed a reporter that is size excluded from re-compartmentalization following nuclear rupture events. This allows for robust detection of factors influencing nuclear integrity in fixed cells. We combined this with an automated image analysis pipeline in a high-content siRNA screen to identify new proteins that both increase and decrease nuclear rupture frequency in cancer cells. Pathway analysis identified an enrichment of nuclear membrane and ER factors in our hits and we demonstrate that one of these, the protein phosphatase CTDNEP1, is required for nuclear stability. Further analysis of known rupture contributors, including a newly developed automated quantitative analysis of nuclear lamina gaps, strongly suggests that CTDNEP1 acts in a new pathway. Our findings provide new insights into the molecular mechanism of nuclear rupture and define a highly adaptable program for rupture analysis that removes a substantial barrier to new discoveries in the field.

## Introduction

Developing mechanistic insight into nuclear membrane rupture is crucial to understanding this potential driver of cancer metastasis and progression, however technological barriers limit our ability to perform broad unbiased screens and define molecular mechanisms. Nuclear membrane rupture is defined as the rapid loss of nucleus compartmentalization in interphase resulting in the mixing of nuclear and cytoplasmic proteins and organelles^1–3^. Ruptures are typically repaired within minutes with limited effects on cell proliferation or viability^4,5^. However, they can cause DNA damage, gene expression changes, and activation of cell invasion and inflammation pathways, especially under conditions of mechanical stress^2,4–9^. In addition, persistent rupture of micronuclear membranes can lead to catastrophic changes in chromosome structure, ongoing chromatin defects, and altered gene expression patterns that are proposed to be initiating events in some tumors and drivers of metastasis^10–15^.

Nuclear membrane rupture occurs in a range of mechanically challenging conditions *in vivo*, including migration through dense extracellular matrices, at the leading edge of tumors, and during nuclear migration in fission yeast, *C. elegans*, and mouse models of laminopathies^4,5,8,16–22^. Changes in nuclear membrane stability underlie pathogen infection, neurodegeneration, autoimmune diseases, cancer progression, and laminopathy pathologies^23,24^, suggesting nuclear envelope dynamics must be strictly regulated for proper cell functioning.

A current model of nuclear membrane rupture is that it occurs in response to an imbalance between force and force resistance at the nuclear envelope^25–27^. In general, membrane rupture occurs when a gap forms in the nuclear lamina matrix, exposing the membrane to cellular forces. This loss of support is frequently followed by membrane blebbing and the simultaneous rupture of the inner and outer nuclear membranes^24,28–31^. Actomyosin forces are the major driver of nuclear rupture in tissue culture and migrating cells. In cultured cells, both increased nuclear compression by perinuclear-actin cables and increased actomyosin contractility have been shown to drive rupture^32–37^. Opposing these forces, increasing the amount of heterochromatin or nuclear lamina proteins increases mechanical resistance and limits nuclear lamina gap frequency, membrane blebs, and rupture^3,30,38–42^. Despite the elucidation of this general model, many questions about rupture remain, including how lamina gaps form, what triggers the transition from membrane expansion to membrane rupture, and how the frequency and size of the rupture affects the cellular consequences of losing nucleus compartmentalization.

The current gold standard for identifying and quantifying nuclear membrane rupture is live-cell imaging of a fluorescent protein tagged with a nuclear localization signal (NLS). Although software exists to automatically quantify ruptures from these data^40^, the sheer size of the data and imaging time required precludes systematic identification of membrane stability regulators using live-cell imaging methods. Several proteins remain at rupture sites for some time after membrane repair, including cGAS, BAF, and lamin A, and have been used to identify ruptures in fixed tissue^21^. However, the extent and duration of their localization can be highly variable and dependent on additional factors^14,43^, limiting their utility as a screening tool. An alternative approach uses mislocalization of large complexes to the nucleus or the cytoplasm to identify cells that have previously lost nuclear integrity. During rupture soluble proteins and small organelles diffuse through the membrane gap and become mislocalized^2,3,26,44^. Chromatin-directed nuclear re-compartmentalization initiates within seconds of the chromatin being exposed to the cytoplasm and proteins containing nucleus and cytoplasmic sorting signals (NLS and NES, respectively) are rapidly relocated^4,33,45,46^. In contrast, organelles and large proteins lacking sorting signals, including Hsp90, 53BP1, and PML bodies, can persist in the wrong compartment for several hours^8,25,44,47,48^.

Building on these observations, we developed a reliable and well-tolerated reporter of nuclear membrane rupture that can be used in fixed cells, called RFP-Cyto. We demonstrate that mis-localization of RFP-Cyto to the nucleus is a rapid, precise, and durable marker of nuclear membrane rupture. We describe an automated analysis pipeline to quantify rupture frequency in fixed cell populations using this reporter and combine it with siRNA screening to discover 22 new regulators of rupture in a high-throughput and unbiased manner. Co-analysis of GFP-NLS localization allowed us to further expand the rupture phenotypes under investigation to include proteins that regulate membrane repair or single cell rupture frequency and identified the kinase STK11 as an additional stability factor. We apply a suite of imaging and analysis tools to validate the nuclear lipin phosphatase CTDNEP1 as a significant rupture inhibitor and delineate its effects on cellular structures associated with nuclear stability. We demonstrate that CTDNEP1 loss does not substantially alter nuclear lamina gap frequency, nuclear pore density, or nuclear confinement, strongly suggesting that it represents a new mechanism of membrane stabilization. This work represents the first systematic identification of nuclear membrane rupture and repair factors in human cells and defines a comprehensive set of tools to rapidly identify factors acting in new pathways. Our tools substantially reduce the barrier to identifying cellular changes that impact membrane stability and pave the way for future genome-wide analyses.

## Results

### RFP-Cyto is size excluded from re-compartmentalization following nuclear rupture

To develop a fixed cell reporter for interphase nuclear rupture, we took advantage of the fact that large objects and proteins lacking a nuclear/cytoplasm sorting signal remain mi-localized after nuclear membrane rupture and repair. We found that a construct containing luciferase tagged with two RFPs was the smallest one (118 kD) which showed exclusive cytoplasm localization during interphase (RFP-Cyto, Fig. 1A, Fig. S1A).We optimized for a minimally sized reporter to increase the likelihood that small, short ruptures would result in reporter mislocalization to the nucleus (Movie S4). In contrast to proteins tagged with an NLS (*e*.*g*. GFP-Nuc), we expected RFP-Cyto would remain mislocalized after membrane repair (Fig. 1A-B).

**Figure 1.**
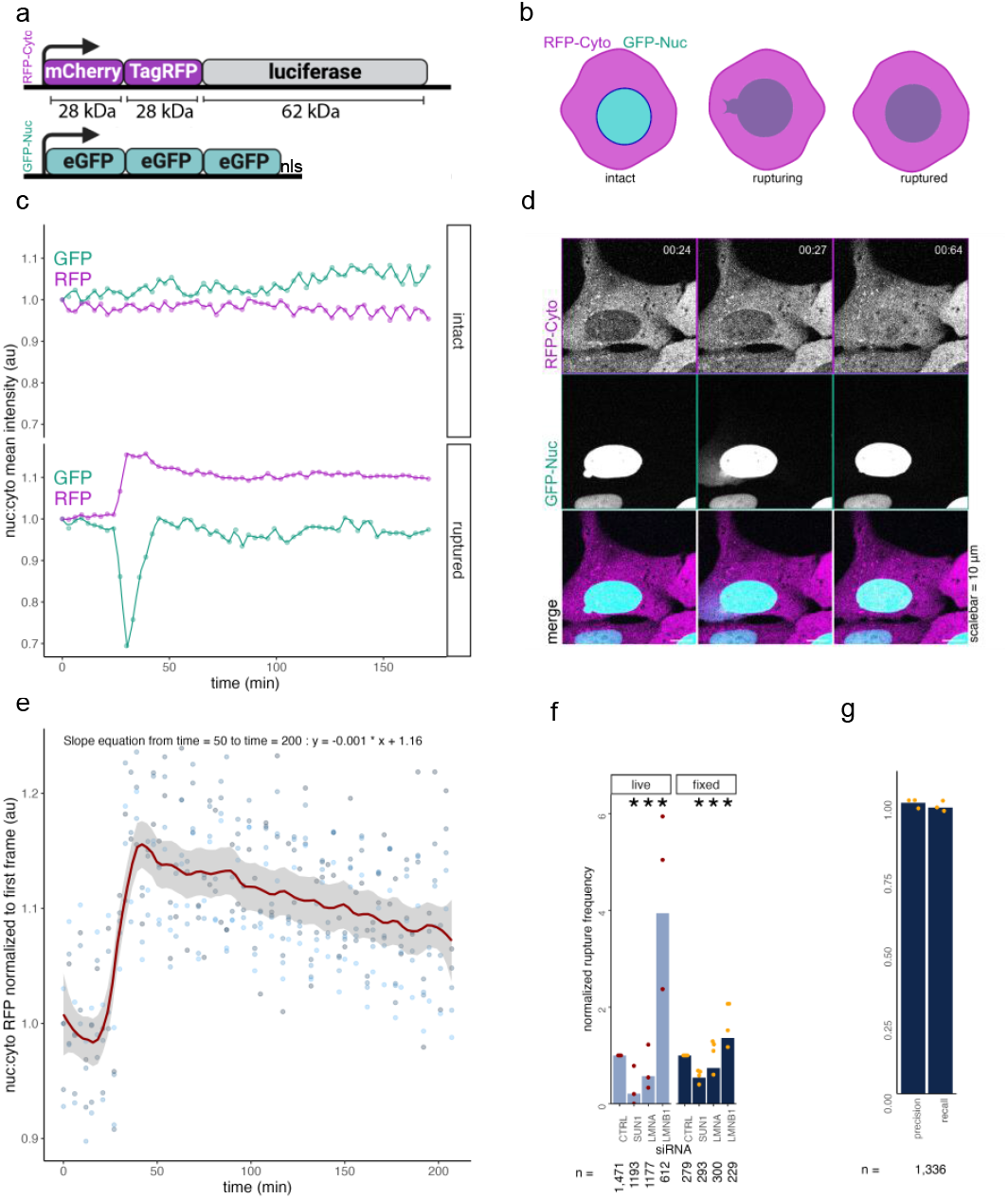
RFP-Cyto is size excluded from re-compartmentalization following nuclear rupture, enabling fixed cell analysis. **a.** Schematic of RFP-Cyto and GFP-Nuc reporters **b**. Schematic of rupturing cells with incorporated reporters **c**. Representative traces of intact and rupturing cells **d**. Representative image panel of cells before, during, and after rupture **e**. Linear model of five cells traced before during and after a single rupture with best fit line, loess smoothing = 0.2, standard error shaded in gray, dots indicate individual datapoints **f**. Manual rupture frequency analysis for live and fixed cells based on RFP-Cyto **g**. Precision and recall for live experiment in (f). Cells: U2OS RuptR, Stats: Table S1, * = p < 0.05

We stably co-expressed RFP-Cyto in U2OS cells expressing GFP-Nuc and an shRNA against LMNB1 and monitored RFP-Cyto localization during nuclear rupture by live-cell imaging. Constitutive expression of anti-LMNB1 shRNAs in U2OS cells, which leads to only a modest reduction in protein levels, substantially increases nuclear rupture frequency and allows for sensitive identification of *in vivo* relevant mechanisms^35,46,49^. We imaged this cell line, U2OS RuptR, for 24 hours with a pass time of 3 minutes and analyzed both GFP and RFP mean nuclear intensities during rupture events. As expected, RFP-Cyto intensity increased in the nucleus after rupture, indicated by a decrease in GFP-Nuc signal, and remained mis-localized after membrane repair, indicated by GFP-Nuc re-compartmentalization. In the absence of nuclear rupture, no change in RFP-Cyto nuclear intensity was observed (Fig. 1C-D, Movie S1). RFP-Cyto persisted in the nucleus for at least 150 minutes after rupture, and nuclei with one or more ruptures often maintained elevated nuclear RFP to the end of imaging (Fig. 1E, Fig. S1C). Importantly, RFP-Cyto was not mislocalized to post-mitotic nuclei, showing no fluorescence soon after nuclear envelope assembly (max GFP-Nuc nuclear intensity)^50,51^ (Fig. S1B, Movie S5).

To determine the accuracy and sensitivity of RFP-Cyto as an interphase rupture reporter, we quantified the proportion of nuclei with nuclear RFP-Cyto in U2OS RuptR cells depleted of lamin B1, lamin A/C, or Sun1 by live or fixed cell imaging and analysis (Fig. S1D). Transfection of siRNAs against LMNB1 in this cell line substantially increases protein loss and rupture frequency^38^. As expected, lamin B1 loss increased rupture frequency and Sun1 loss suppressed rupture frequency by live-cell analysis of GFP-Nuc^35,38^ (Fig. 1F, left). Although lamin A/C depletion can increase rupture frequency^2,40^, we did not see a consistent increase upon lamin A/C loss, in line with recent results from other systems^31,37^. Similar results were observed when ruptures were analyzed using RFP-Cyto in fixed cells (Fig. 1F, right). Comparison of RFP-Cyto and GFP-Nuc mislocalization during live-cell imaging demonstrated that RFP-Cyto nuclear localization accurately predicted nuclear rupture events, with a pooled precision of 99% and recall of 99% (Fig. 1G). Rare ruptures missed by RFP-Cyto tended to be short and in cells with low RFP-Cyto expression (Movie S2-S4). These data demonstrate that analysis of fixed U2OS RuptR cells provides accurate information about relative rupture frequencies and with a high degree of sensitivity.

### An automated analysis pipeline accurately detects alterations to nuclear stability in U2OS RuptR cells

To enable high throughput analysis of RFP-Cyto localization and intensity, we developed a segmentation strategy in CellProfiler^52^ coupled with a set of post-segmentation intensity and morphological filters in R to remove several classes of contaminating objects. This workflow was developed on U2OS RuptR cells transfected with siRNAs, arrested in S phase for 24 h to enhance rupture frequency and reduce mitotic false positives^35^, fixed, and labeled with Hoechst to mimic screening conditions (Fig. 2A). Nuclei and cells were segmented in CellProfiler based on Hoechst and RFP-Cyto signal, respectively, and morphological parameters were analyzed (Fig. S2A, Fig. 2B-C). Objects were then shrunk to limit overlapping signal or segmentation errors and RFP, GFP and Hoechst mean intensity in the cytoplasm and nucleus were measured (Fig.

**Figure 2.**
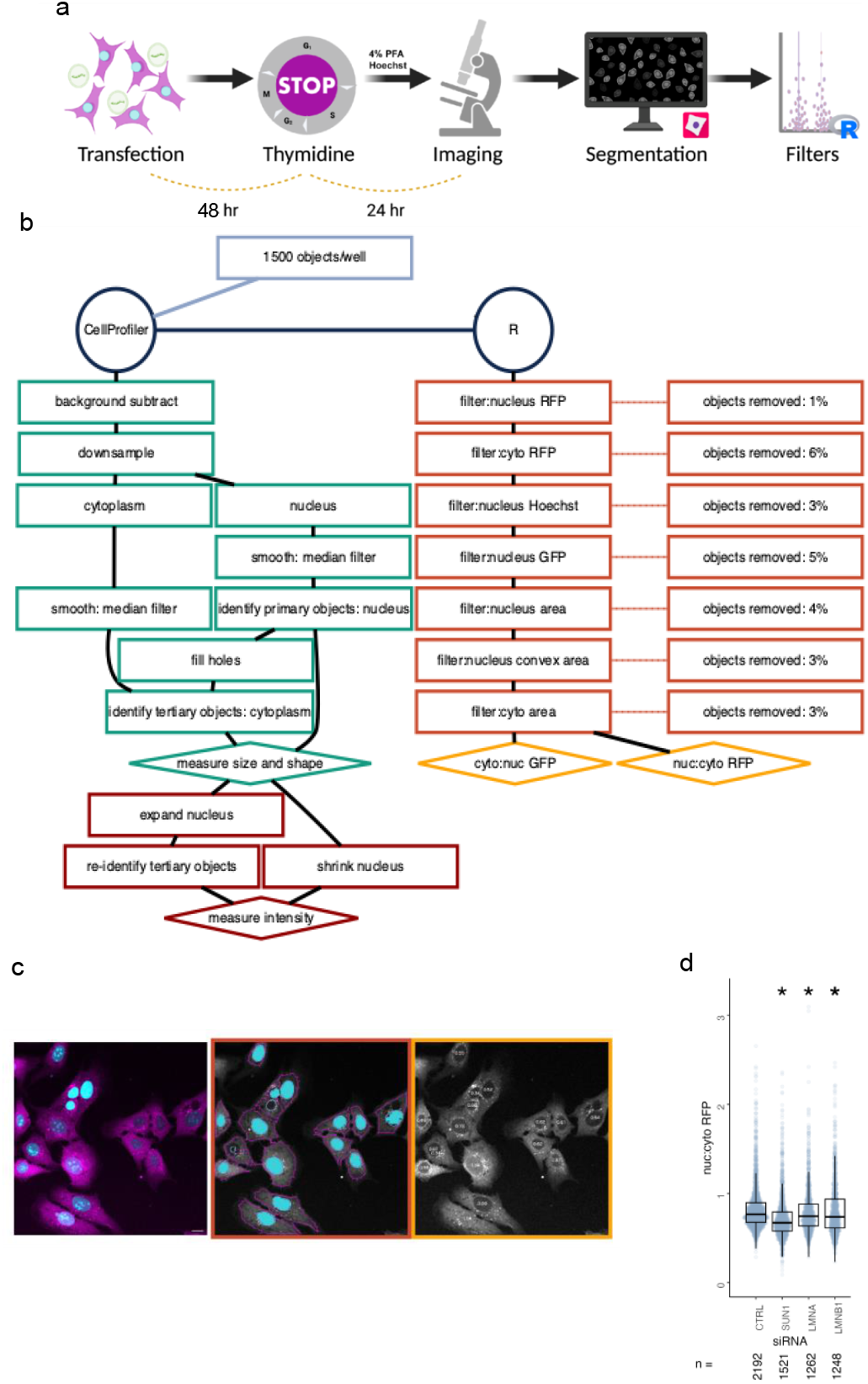
An automated bioinformatics pipeline allows accurate fixed cell analysis of rupturing cells. **a.** Schematic of experimental pipeline **b**. Detailed view of data processing steps **c**. Representative images of one field of view left: merge, center: false colored nuclei passing filtration, right: overlay with nuc:cyto RFP values **d**. Nuc:cyto mean intensity RFP for cell populations treated with siRNA as listed. Cells: U2OS RuptR, Stats: Table S2, * = p < 0.05.

2B, Fig. S2A). Intensity and morphology filters were applied post-processing to eliminate data from cells with low RFP-Cyto expression, mis-segmented objects, mitotic or dead cells, and out of focus cells. Mitotic cells were removed by setting a maximum Hoechst intensity threshold, coupled with a maximum solidity threshold, and out of focus cells were reduced by setting a minimum Hoechst intensity threshold. Cells undergoing cell death were limited by setting minimum nuclear size and solidity thresholds (Fig. S2B). Filter thresholds were set globally for intensity measurements and per condition for morphology measurements to accommodate expected variations in nucleus shape and size during screening (Fig. 2B, Fig. S2A). After filtering, over 75% of cells were retained and analyzed for RFP-Cyto localization by calculating the nucleus to cytoplasm (nuc:cyto) mean intensity ratio.

To validate this pipeline, U2OS RuptR cells were reverse transfected with pooled siRNAs targeting LMNB1, SUN1, and LMNA. Following knockdown for 48 h, cells were arrested in S phase for a further 24 h, fixed, and labeled with Hoechst before imaging. An analysis of the distribution of the nuc:cyto RFP ratios for siRNA pools against LMNB1 and SUN1 as positive controls and a non-targeting siRNA pool as a negative control identified significant (Wilcoxon rank-sum test) differences between these conditions (Fig. 2D). These results demonstrated that our screen design was sufficiently sensitive to identify biologically relevant changes in rupture frequency.

### A high content screen identifies new factors influencing nuclear integrity

To identify new regulators of nuclear membrane rupture, we applied this pipeline to a plate-based siRNA screen targeting known categories of nuclear membrane stability regulators (49%) and randomly selected genes (51%) (Table S9). Targeted categories, all of which were enriched overall within the screen (Fig. S4A), included chromatin organization^30^, tumor suppressors^53–55^, nuclear envelope localized^2,3,31,40,56^, ER/nuclear membrane structure^57^, and cytoskeleton^35,37,58,59^. Images from at least three technical replicates were analyzed and the nuc:cyto RFP mean intensity ratio was calculated on cells passing all filters and the proportion of cells with ruptured nuclei (“RFP+”) were calculated for each condition. The threshold for RFP+ cells was defined as a nuc:cyto RFP ratio greater than one standard deviation above the median of the three CTRL siRNA wells present in each technical replicate. To determine the robustness of our image analysis pipeline, images from siRNA conditions with statistically significant changes in nucleus size and solidity (Fig. S3A) or in median RFP-Cyto intensity were manually evaluated for segmentation and RFP+ errors. Our automated analysis and filtering was robust to each of these parameters, with a minimum precision of 80% and a recall of 94% and a consistently high frequency of proper cell and nucleus segmentation (Fig. S3B-C). Visual inspection also led to the exclusion of PSMC1 knockdown, an outlier in fluorescence intensity, from further analysis due to excessive cell death, consistent with its function as an essential proteosome component^60^ (Fig. S3D).

Our screen successfully identified 22 genes as hits (24% of screen), with 14 (15%) decreasing rupture frequency when depleted and 8 (8.6%) increasing it, based an adjusted p-value cut-off of 0.05 (Barnard’s test, Bonferroni correction for multiple testing) and a log_2_ fold-change cut-off of 0.3 (Fig. 3A, Fig. S3E, Table S3). To validate our screen results, we selected seven hits to re-analyze using single siRNAs. Nuc:cyto RFP intensity ratio distributions were quantified using the same experimental design as the screen. We found 6/7 targets significantly altered nuc:cyto RFP (Wilcoxon rank-sum test)(Fig. 3B), demonstrating that our screen allows unbiased identification of nuclear membrane stability factors with a high degree of accuracy.

**Figure 3.**
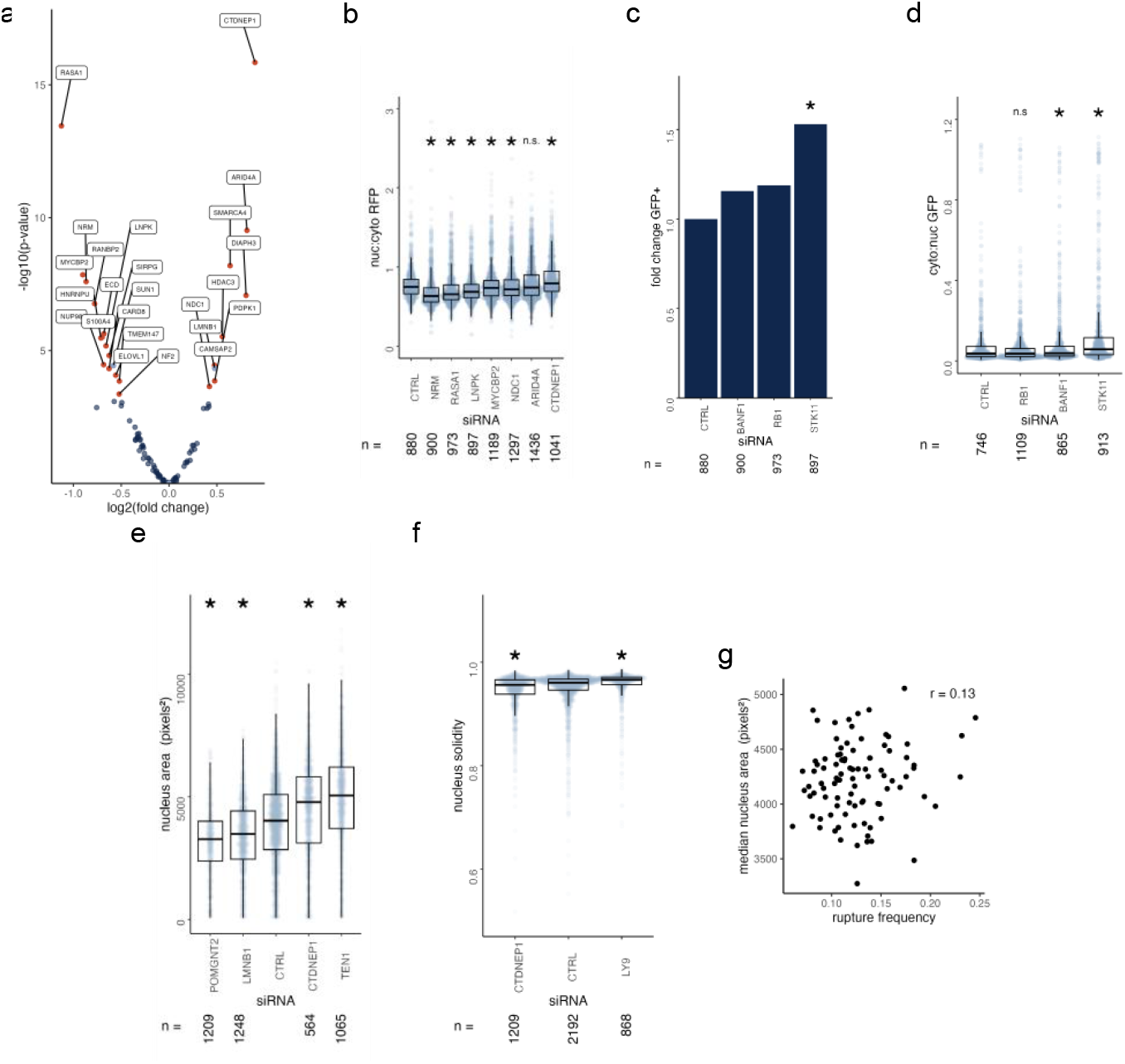
A high content siRNA screen accurately identifies factors influencing nuclear integrity and morphology. **a.** Volcano plot indicating siRNAs that significantly influenced nuclear integrity determined by nuc:cyto RFP ratio over threshold, labels indicate Bonferroni adjusted Barnards test p < 0.05 **b**. Distribution of individual cells transfected with single siRNAs for selected high and low RFP-Cyto screen hits **c**. Fold change GFP+ results for siRNAs with > 175 RFP+ cells showing increase in GFP-Nuc loss of compartmentalization **d**. Distribution of individual cells transfected with single siRNAs for selected GFP-Nuc screen hits **e**. Nuclear area distribution for selected screen hits **f**. Nucleus solidity distributions for selected screen hits **g**. Factors influencing nucleus area did not strongly correlate with higher rupture frequency (Pearson). Cells: U2OS RuptR, Stats: Table S3-S8, * = p < 0.05.

We further extended our screen to quantify conditions increasing the proportion of rupturing, versus ruptured, nuclei (Fig. 1B) to identify proteins that inhibited membrane repair and/or increased single cell rupture frequency. The proportion of ruptured cells with GFP-Nuc mislocalized to the cytoplasm, defined as a cyto:nuc GFP mean intensity greater than one standard deviation from the median of the three siCTRL populations, was calculated for each condition with ≥ 175 ruptured nuclei cells identified per replicate. As a positive control, we depleted BANF1(BAF), which is required for efficient membrane repair^45,46^. Of the 26 screened siRNAs with sufficient rupture events, only depletion of RB1 and STK11 increased the frequency of RFP+/GFP+ double positive cells at or above the level of BANF1. To validate these hits, single siRNAs against BANF1, RB1, and STK11 were transfected into U2OS RuptR cells and the distributions of cyto:nuc GFP ratios were analyzed (Wilcoxon rank-sum test). Both BANF1 and STK11 depletion significantly increased the median ratio, strongly suggesting that STK11, a cancer-associated kinase that acts as a master regulator of cell growth and metabolism^61^, has additional functions in maintaining nuclear integrity.

Changes in nucleus morphology can be strongly correlated with changes in nucleus stability^62^. Our analysis of nuclear area and nucleus solidity identified five conditions that significantly altered nuclear size or lobulation, including LMNB1, a known nuclear growth factor^63,64^, and CTDNEP1, a known nuclear size restriction factor^65^ (Fig. 3E-F). Despite having opposite effects on nuclear size, both CTDNEP1 and LMNB1 depletion increased nuclear rupture, and Pearson correlation analysis found no significant relationship between nucleus rupture and median nucleus area in the screened genes (Fig. 3G).

To determine whether screen hits were enriched in specific cellular pathways, we performed GO (gene ontology) analysis with hypergeometric enrichment statistics to correct for the bias towards nuclear factors in our gene set (Fig. S4A). Overall, our hitlist was only slightly enriched in targeted compared to randomly selected genes (Fig. S4B). GO term analysis identified a high enrichment in predicted processes, including cytoskeletal organization and nuclear protein complexes, as well as unexpected processes and components, including tissue development and the outer nuclear membrane /ER network (Fig. 4). Although membrane shaping proteins were a targeted gene class, their enrichment over proteins in the inner nuclear membrane and nuclear pore strongly suggests that these factors are a new key regulator of nuclear stability.

**Figure 4.**
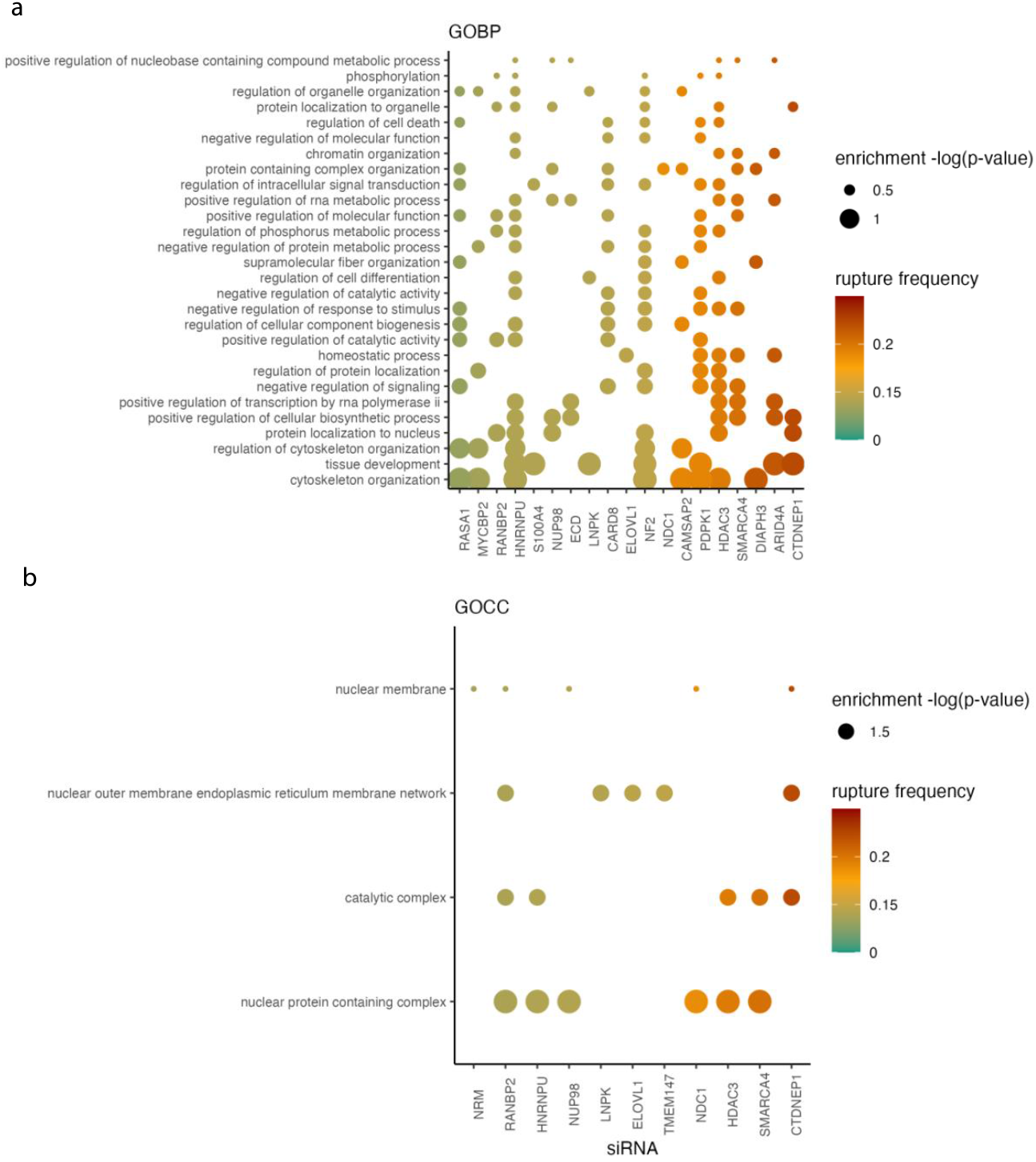
Hypergeometric enrichment identifies nuclear and cellular pathways influencing nuclear integrity a,b. Hypergeometric enrichment of gene ontology terms (**a**: GOBP, **b**: GOCC) for siRNAs with significant RFP+ screen results, rupture frequency indicated by color, p-value indicated by size. Stats: Table S10-11.

### CTDNEP1 maintains nuclear membrane stability in interphase through a new pathway

The unanticipated enrichment of ER/nuclear membrane factors in our screen hits prompted us to further characterize the function of one of these proteins, CTDNEP1, in nuclear membrane stability. CTDNEP1 is a protein phosphatase that localizes to the nuclear membrane and regulates lipid composition through activation of the phosphatidic acid phosphatase, lipin1^66^. CTDNEP1 caused the largest increase in nuclear rupture upon depletion in our screen (Fig. 3A) and additional validation with a second siRNA and qRT-PCR confirmed the depletion of CTDNEP1 and the specificity of the nuclear rupture phenotype for CTDNEP1 loss (Fig. S5A-B). As a positive control for CTDNEP1 depletion, we examined ER area and nuclear size and found that both were increased, as expected, in our cells^65,67^ (Fig. 3E, Fig. S5C-E). Live-cell imaging analysis of CTDNEP1 siRNA depletion was performed in U2OS shRNA-LmnB1 and HeLa cells expressing GFP-Nuc and caused a significant increase in rupture frequency in both cell lines, further validating our fixed-cell analysis and demonstrating that this effect is not cell type specific nor dependent on lamin B1 depletion (Fig. 5A, Movie 6). CTDNEP1 loss can delay nuclear membrane resealing after division^65^, therefore we analyzed rupture duration by live-cell imaging of GFP-Nuc in U2OS RuptR cells. We found no statistical difference between control and CTDNEP1 depletion (Fig. 5B), consistent with the results of our fixed-cell GFP-Nuc screen, suggesting that a requirement for CTDNEP1 in membrane closure is context dependent. CTDNEP1 loss did not significantly increase micronucleus rupture (Fig. 5C), strongly suggesting that CTDNEP1 is a new broad-acting inhibitor of membrane rupture specific to the main nucleus.

**Figure 5.**
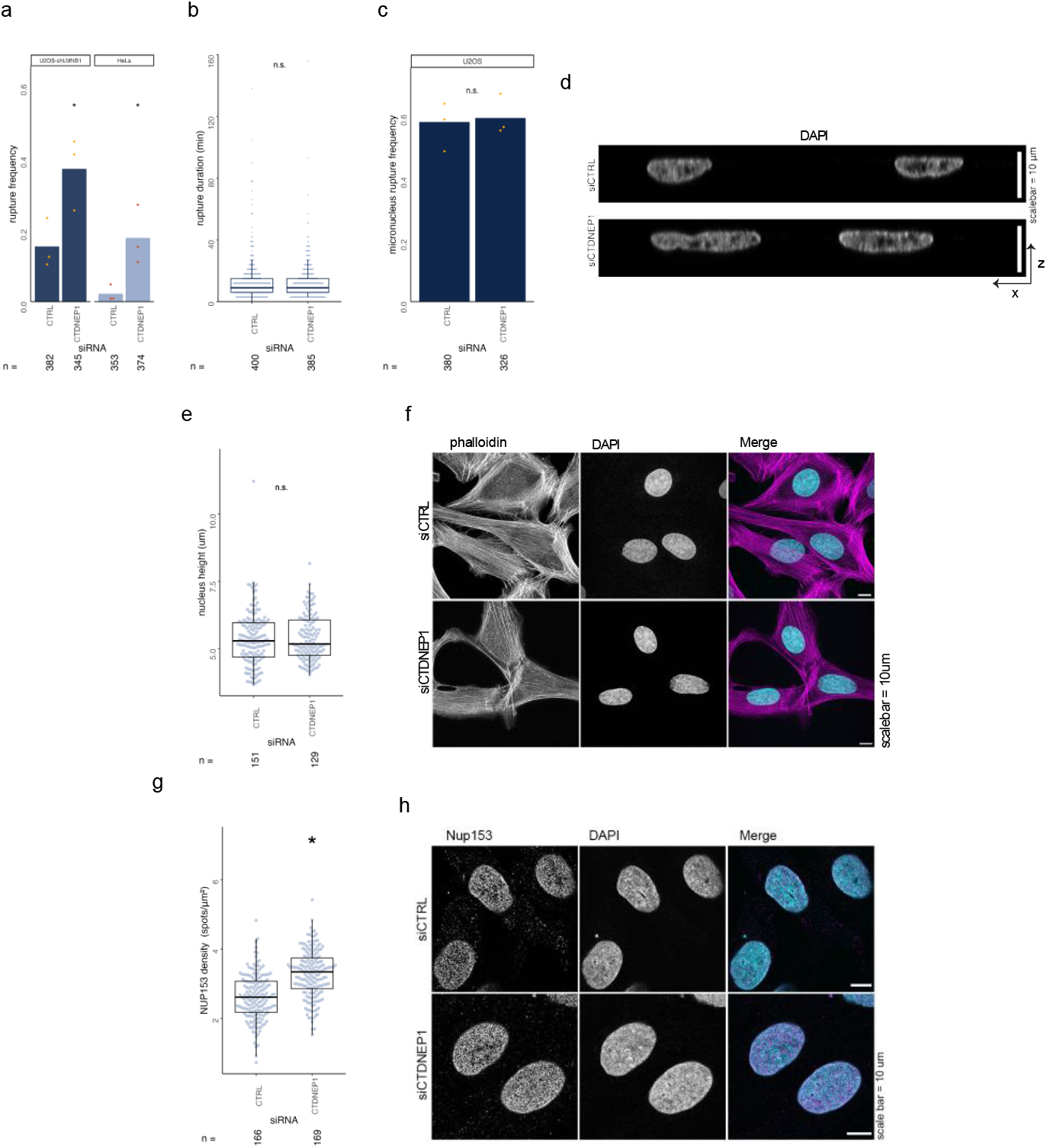
CTDNEP1 loss increases rupture frequency in the absence of nuclear pore or confinement defects. **a.** Single CTDNEP1 siRNA increases rupture frequency in U2OS-shLMNB1 RFP-NLS and HeLa GFP-Nuc cells by live-cell imaging. * = p < 0.05, Barnards test. **b**. CTDNEP1 loss does not increase rupture duration by live-cell imaging **c**. Micronucleus rupture frequency in siRNA treated U2OS GFP-Nuc cells **d**. Representative images showing nucleus height (DAPI) in siRNA treated cells **e**. Quantification of nuclear height depicted in (d) shows no significant change. **f**. Representative images showing actin (phalloidin) in siRNA treated U2OS GFP-Nuc cells **g**. NUP153 nuclear pore quantification showing increase in density following CTDNEP1 depletion in U2OS GFP-Nuc cells. **h**. Representative images of NUP153 foci, single section. * = p < 0.05, KS test. Cells: U2OS shLMNB1 2xRFP-NLS unless indicated, Stats: Table S14-S17.

We next sought to determine the mechanism by which CTDNEP1 stabilizes the nuclear membrane. Analysis of nuclear height found no significant difference between CTDNEP1 and control depleted cells (Fig. 5D-E), and no changes in perinuclear actin organization (Fig. 5F), strongly suggesting that CTDNEP1 loss does not increase nucleus confinement. We next quantified nuclear pore density as changes in nuclear transport have been linked to increased nuclear membrane tension^36^ and CTDNEP1 has functions in nuclear pore formation^68^. Depletion of CTDNEP1 caused a slight, but significant, increase in nuclear pore density (Fig. 5G-H), strongly suggesting that CTDNEP1 loss is unlikely to increase nuclear rupture via nuclear transport defects.

A major predictor of nuclear membrane rupture in both nuclei and micronuclei is the presence of nuclear lamina gaps. To robustly and sensitively quantify nuclear lamina gaps, we developed an application that uses distance transformation on binarized images of DAPI and a nuclear lamina protein, here lamin A, to define the presence, number, and nuclear lamina gaps per nucleus, as well as the relative curvature of the nucleus at gap locations (Fig. 6A-B). To validate this method, we compared nuclear lamina gaps in S-phase arrested U2OS cells with or without shRNAs against LMNB1, which significantly increases gap number^35^. As expected, LMNB1 depletion significantly increased the number of nuclei with nuclear lamina gaps and the gap number per nucleus (Fig. 6C-D, CTRL siRNA), but not lamina gap size (Fig. 6E). Lamina gaps are frequently located at the highly curved poles of the nucleus^25,69^. Our analysis confirmed that gaps are enriched on curved surfaces in both cell lines when few gaps are present (1-3), but also showed an increased localization (curvature = 0 um^-1^) to flat nuclear surfaces as the gap number increases (Fig. 6F).

**Figure 6.**
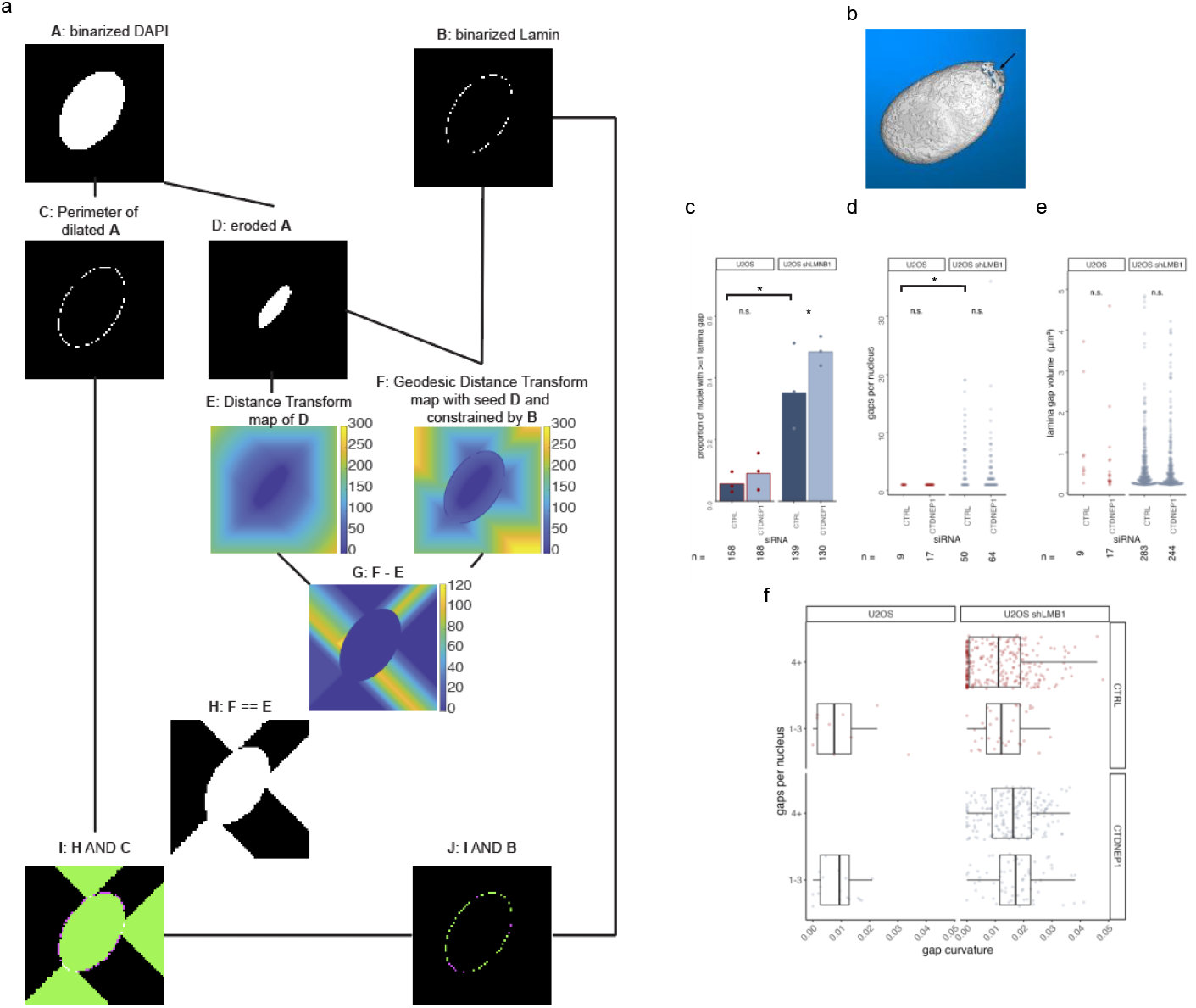
Automated quantification of nuclear lamina gaps identifies minimal increase in lamina gap frequency from CTDNEP1 loss. **a**. Overview of the image processing steps used to identify and characterize gaps in the nuclear lamina, full description available in methods **b**. Rendering of nucleus from the lamina gap analysis tool with one gap indicated by arrow **c-e**. Characterization of lamina gaps showing proportion of nuclei with at least one gap (c), gaps per nucleus where at least one gap was identified (d), and gap size (e) in U2OS GFP-Nuc or U2OS shLMNB1 2xRFP-NLS cells treated with siRNAs. * = p < 0.05, Barnard’s test (c), or Kruskal-Wallis test (d-e). For (e) pKW = ns. **f**. Gaps per nucleus plotted against gap curvature for siRNA treated shows gaps on flat surfaces limited to high gap number and gaps are enriched on curved surfaces in CTDNEP1 depleted cells. Stats: Table S19-S20.

Together these data strongly suggest that depletion of lamin B1 alters the frequency, but not the physical characteristics, of nuclear lamina gaps and validate our automated approach. We next applied this analysis to cells depleted of CTDNEP1 and found a small increase in the proportion of nuclei with nuclear lamina gaps and no increase in gap number or size (Fig. 6C-E). The limited and inconsistent increase in cells with nuclear lamina gaps, and the lack of change in gap frequency, unlike after lamin B1 depletion, strongly suggests that increased lamina disruption is unlikely to fully explain the ∼2.5 fold increase in rupture frequency upon CTDNEP1 depletion (Fig. 6A). Unexpectedly, CTDNEP1 loss reduced the frequency of gaps on flat nuclear surfaces (Fig. 6F). A partial explanation may be the increased 2D curvature of CTDNEP1 depleted nuclei (Fig. 3F, S3A), but orthosection and height analyses indicate that CTDNEP1 loss does not reduce the amount of flat nuclear surfaces (Fig. 5D-E), suggesting that another mechanism is driving lamina gaps clustering at the poles. Together, our results strongly suggest that CTDNEP1 acts in part through exacerbating existing nuclear lamina defects but also through a new mechanism of nuclear membrane rupture.

## Discussion

We have developed a new screening pipeline, based on automated image analysis of a nuclear-excluded reporter, that enables rapid identification of conditions that impact nuclear stability without the need for live-cell imaging. This system allows us for the first time to define novel cellular regulators of nuclear membrane rupture in an unbiased manner. In addition to paving the way for future large-scale screens, our fixed-cell technique will accelerate analysis of molecular mechanism experiments in this field and enable characterization of nuclear rupture in samples that are not amenable to long-term live-cell imaging, including rapidly moving cancer cells and 3D organoids. We expect this technology will be adaptable to any cell type that permits exogenous protein expression and has a cytoplasm compartment that can be segmented. In addition, this study defines a suite of automated image analysis methods, including a new open-source application for nuclear lamina gap quantification, that can be employed in a secondary screen to place hits in functional pathways based on their effects on nuclear structure. By creating standardized methods for these analyses, our tools will also facilitate comparison of nuclear structures across systems and enable detailed molecular analysis of mechanisms that disrupt the nuclear lamina. Overall, our new reporter system lowers the barrier of entry for biologists in any field to evaluate the contribution of nuclear membrane rupture to their phenotype of interest and quantitatively identify changes in contributing cellular structures to accelerate mechanistic dissection.

Nuclear lamina gaps have been observed in both developmental and disease contexts in a broad group of species^24,70^. They have been the subject of much modeling and analysis^33,69,71–73^, yet fundamental questions about them remain, including how they form, how long they persist, and whether they can be repaired. One limiting factor in generating these models is the lack of large quantitative datasets. The nuclear lamina analysis application developed in this study overcomes this challenge by providing robust and rapid quantification of many gap parameters, including number, size, solidity, and nuclear curvature from 3D fluorescence images. Additional nuclear morphometric parameters are included in the output including height, volume, surface area, and label intensity. Our initial analysis of the number, size, location of lamina gaps in U2OS cells with and without lamin B1 depletion, strongly suggests that gaps form and expand by the same mechanism in both conditions, but that the initial barrier to forming a gap is lowered by decreased lamin B1. Similar analyses of other conditions increasing nuclear lamina gap frequency may further stratify these phenotypes and reveal new mechanisms.

Our initial siRNA screen against targeted and randomly selected genes had a high hit and validation rate for both groups, strongly suggesting that nuclear rupture occurs in many more conditions than are currently known. In addition, our results suggest that changes in nuclear membrane stability is a sensitive assay to detect alterations in nuclear or cytoskeletal structure and that our system could be a valuable tool for uncovering new regulators of both. This screen also identified proteins that promote membrane nuclear membrane rupture in cancer cells. Future analysis of how these proteins act will provide new insight into how cells can adapt to high rupture rates and their depletion will enable critical experiments defining the specific consequences of membrane rupture in cells with disrupted nuclear structures.

A major gap in our understanding of nuclear membrane rupture is how the membrane itself responds to mechanical stress. Several lipid-based mechanisms have been identified that could relieve nuclear membrane tension and prevent nucleus rupture^18,19,74–80^, but whether these pathways are active during nuclear stress in mammalian cells is unknown. Our results support a role for membrane structural proteins in determining rupture frequency. Of the proteins driving GO:CC enrichment in nuclear/ER membrane categories, the majority regulate the physical structure of the membranes, including CTDNEP1, LNPK, Ndc1, and EVOLV1^67,81–89^.

CTDNEP1 regulates nuclear membrane amount and lipid composition through activation of the phosphatidic acid phosphatase, lipin1^67,74,83,84,86,90^, which has downstream effects on chromosome segregation, post-mitotic nuclear assembly, nuclear pore insertion, and protein stability^65,67,80,87,91^. CTDNEP1 may also regulate major signaling pathways, including myc, BMP and WNT, that can affect actin organization through lipin-independent mechanisms^92–98^. Our analysis of nuclear features after CTDNEP1 depletion demonstrates that the increase in membrane rupture is due to a new mechanism not linked to nuclear lamina organization, actin bundle organization, or nuclear pore insertion defects. Defining the molecular mechanism of how CTDNEP1 regulates nuclear membrane stability, and its dependence on lipin1, could provide new information on how lipid composition contributes to nuclear membrane stress or how signaling pathways feedback to nuclear envelope structure.

In this study we developed several new tools that enable comprehensive identification of proteins and conditions that regulate nuclear membrane stability and their putative mechanism of action. This technology enabled us to identify 22 potential new rupture factors from a small-scale screen, and to demonstrate that one of these new factors, CTDNEP1, provides critical support for the nuclear membrane through a new mechanism potentially centered on lipid metabolism. This pipeline can be easily modified to identify specific rupture characteristics, including correlation with membrane blebs, or the integrity of other compartments, including micronuclei. Overall, the ability to rapidly quantify changes in nuclear rupture frequency using standard confocal microscopy will accelerate our understanding of the molecular mechanisms regulating nuclear envelope structure and dynamics and its consequences during development and disease.

## Supporting information

Table of Contents for Supplementary Movies and Tables

## Acknowledgements

E.M.H, A.L.G., and A.Y. were supported by the National Institutes of Health grant R35GM124766. J.J. was supported by the CMB training grant T32 GM007270. This work was supported by the Cellular Imaging and Flow Cytometry Shared Resources of the Fred Hutch/University of Washington Cancer Consortium (P30 CA015704) and Fred Hutch Scientific Computing, NIH grants S10-OD-020069 and S10-OD-028685. Schematics were generated with BioRender.com. This paper was typeset with the bioRxiv word template by @Chrelli: www.github.com/chrelli/bioRxiv-word-template

## Author contributions

ALG: conceptualization, data curation, formal analysis, investigation, methodology, project administration, validation, visualization, writing – original draft, review and editing

AIY: data curation, formal analysis, investigation, methodology, visualization, writing – original draft, review and editing

JD: data curation, methodology, writing – review and editing. JJ: formal analysis, writing – review

EMH: conceptualization, formal analysis, funding acquisition, investigation, methodology, project administration, resources, supervision, visualization, writing – original draft, review & editing

## Materials and Methods

### Cell lines and transfection

U2OS (ATCC:HTB-96) and HeLa (ATCC:CCL-2) cells were grown in 1x DMEM (GIBCO) plus 10% FBS (Sigma-Aldrich), 1% Penicillin-Streptomycin (GIBCO) at 10% CO_2_ and 5% CO_2_ respectively. U2OS shLMNB1 2xRFP-NLS, U2OS 3xGFP-NLS, and HeLa 3xGFP-NLS cells were characterized previously^46^. siRNAs were transfected using siLentfect (Bio-Rad Laboratories) according to the manufacturer’s instructions and cells were analyzed at least 72 h post-transfection. RFP-Cyto was introduced by stable transduction into U2OS shLMNB1 GFP-NLS cells (RuptR) characterized previously^35^. U2OS cells were arrested in S phase by a single addition of 2 mM thymidine (Sigma-Aldrich), diluted fresh in PBS, 24 h before and during imaging. HeLa cells were arrested in S phase by a single addition of 2 mM hydroxyurea (EMD Millipore) diluted in water.

### Plasmids and siRNAs

1x and 2xRFP-luciferase constructs were made in pEGFP-C1 backbone by replacement of EGFP with TagRFP, then PCR and ligation of luciferase (1xRFP-luciferase), then PCR and ligation of mCherry with C-terminal sequence N-terminal to TagRFP (2xRFP-luciferase). pLVXE-Blast::mCherry-TagRFP-Luciferase (RFP-cyto) was constructed by PCR and ligation of mCherry-TagRFP-Luciferase from 2xRFP-luciferase into a pLVXTight lentiviral vector (Clontech) previously modified to contain the EF1a promoter and blasticidin resistance.

Fig. 1F siRNA validation: siRNAs against CTRL (non-targeting, cat#: D-001810-01-05), LMNA (5’-GGUGGUGACGAUCUGGGCUuu-3’), LMNB1 (5’-CGCGCUUGGUAGAGGUGGAUUuu-3’), and SUN1 (5’-ACCAGGUGCCUUCGAAAuu-3’) purchased from Horizon Discovery. Cells were left in 35 nM transfection medium for 48 h and assessed after 72 h.

siRNA screen: ON-TARGETplus SmartPool siRNAs were purchased in a 96-well plate from Horizon Discovery according to Table S9. U2OS cells were left in 35 nM transfection media 48 h and assessed after 72 h.

Single siRNA screen validation: ON-TARGETplus siRNAs were purchased from Horizon Discovery. siRNAs: CTRL (cat#: D-001810-01-05), LNPK (cat#: J-023148-09), RASA1 (cat#: J-005276-07), NRM (cat#: J-012779-22), ARID4A (cat#: J-003949-05), NDC1 (5’-CUGCACCACAGUAUUUAUAUU-3’), CTDNEP1 (cat#: J-017869-09), MYCBP2 (cat#: J-006951-05), BANF1 (5-AGUUUCUGGUGCUAAAGAAuu-3’), RB1 (5’-GAACAGGAGUGCACGGAUA-3’), STK11 (5’-UGAUGUGGUGCCGUACUU-3’). Individual siRNAs were transfected at 35 nM into U2OS shLMNB1 cells for 48 h and assessed after 72 h.

Fig. 5,6, S5: siRNAs were purchased from Horizon Discovery for CTDNEP1-9 (cat#: J-017869-09), and CTDNEP1-10 (cat#: J-017869-10). Control siRNAs are same as single siRNA screen validation. siRNAs were transfected at 50 nM in transfection medium and left on overnight (U2OS), or 5 h (HeLa).

### Fixed Cell Imaging

Samples fixed 45 m in 4% paraformaldehyde (Electron Microscopy Sciences) in PBS were stained 20 m with 1μg/mL Hoechst (Life Technologies) and imaged as single confocal slices with a 40x/0.75 Plan Apo objective on a Leica DMi8 with a Yokogawa CSU spinning disc, Andor Borealis illumination, and an ASI automated stage with Piezo Z-axis. Images were captured with an Andor iXon Ultra 888 EMCCD camera using MetaMorph software (version 7.10.4; Molecular Devices) equipped with a plate acquisition journal. Acquisition parameters were set to capture 30 images per well. Images were corrected for background signal by mask image subtraction prior to intensity measurements and image displays exhibit rescaled intensity as noted in the CellProfiler 4.2.4 pipeline or modified for brightness/contrast in FIJI^99^. For fixed cell precision/recall analysis, cells were manually assessed for RFP-Cyto in the nucleus and results compared to the RFP+ determinations made by the automated pipeline.

### Time-lapse imaging

Time-lapse images are single confocal slices captured with Leica DMi8 spinning disk microscope and 40x/0.75 Plan Apo objective, as described in Fixed Cell Imaging, at 3 m intervals over 18-24 h. Image sequences were adjusted for brightness and contrast using Fiji. For nuc:cyto mean intensity over time cell traces, nucleus and cytoplasm were manually segmented in Fiji for each time point. For live precision/recall analysis, true positives were manually determined by rapid loss of GFP-Nuc from the nucleus into the cytoplasm co-incident with rapid increase of RFP-Cyto in the nucleus.

### Immunofluorescence

**Calreticulin:** U2OS cells were grown on glass coverslips and fixed 10 m in methanol at −20°C and rehydrated 10 m in PBS at RT. Coverslips were blocked in 3% BSA in PBS plus 0.4% Triton X-100 for 30 m at RT and incubated in primary antibody rabbit anti–Calreticulin at 1:100 (Cell Signaling Technology) overnight at 4°C. Goat anti rabbit Alexa Fluor 647 (Thermo Fisher Scientific) secondary antibody was diluted in blocking buffer 1:1000 and used for 30 m at RT. Coverslips were briefly incubated in 1 μg/mL DAPI (Roche) in PBS and mounted in Vectashield (Vector Laboratories). **Actin:** U2OS-shLMNB1-RFP-NLS cells were fixed 10 m in 4% PFA. Coverslips were blocked in 3% BSA in PBS plus 0.4% Triton X-100 for 30 min at RT and incubated in Alexa Fluor 488 Phalloidin (Cell Signaling Technology) at 1:1000 for 30 min. **NUP153:** U2OS GFP-NLS cells were fixed 10 m in 4% PFA. Coverslips were blocked in 3% BSA in PBS plus 0.4% Triton X-100 for 30 m at RT and incubated in primary antibody mouse anti-Nup153 (Abcam) at 1:250 for 30 m. Goat anti mouse Alexa Fluor 568 (Thermo Fisher Scientific) secondary antibody diluted in blocking buffer 1:1000 and used for 30 m at RT. **Nuclear lamina:** U2OS-shLMNB1-RFP-NLS cells were fixed 10 m in 4% PFA. Coverslips were blocked in 3% BSA in PBS plus 0.4% Triton X-100 for 30 min at RT and incubated in primary antibody mouse anti-laminA (Sigma Aldrich) at 1:500 overnight at 4 °C. Goat anti rabbit Alexa Fluor 488 (Thermo Fisher Scientific) secondary antibody diluted in blocking buffer 1:1000 and used for 30 m at RT.

### qRT-PCR

Cells were harvested from a confluent well in a 6-well plate with 0.25% Trypsin-EDTA, pelleted, and frozen at -80°C. RNA was extracted using RNeasy Mini Plus kit (Qiagen) following manufacturer’s instruction. cDNA was synthesized using iScript RT Supermix (BioRad). A 5ng/μL cDNA reaction mix was set up according to manufacturer’s instructions (SYBR green MM, Fisher Scientific). Thermal cycling was performed at 50°C for 2 min then 95°C for 10 min then 40 cycles of 95°C for 15s, 60°C for 60 s QuantStudio™ 5 Real-Time PCR System (ThermoFisher). Oligo sequences: 5’-AACAGCCTCAAGATCATCAGC -3’ (GAPDH forward), 5’-AGGAGCCATCCAGACAATGC-3’ (CTDNEP1 forward), 5’-CACCACCTTCTTGATGTCATC-3’ (GAPDH reverse), 5’-TCAGCACGGAACGAACATCA-3’ (CTDNEP1 reverse). Ct values for CTDNEP1 were normalized internally to GAPDH and then to transcript levels in cells treated with non-targeting siRNA.

### Western Blot

72 h after siRNA depletion cells were lysed directly in 1x LDS sample buffer (cat#: NP0007, Thermo Fisher) with β-mercaptoethanol. Proteins were separated in 4-20% precast polyacrylamide gels (cat#: 4561096, Bio-Rad) then transferred to nitrocellulose membrane. Membranes were blocked in 5% milk:TBST then incubated 1 h RT with primary antibodies, washed, then incubated with secondary antibodies as noted below. Treated membranes were imaged on an Odyssey (LI-COR Biosciences), and bands were quantified by background subtracting the integrated density and normalizing to tubulin bands using Fiji/ImageJ. Primary antibodies: LMNA (1:1000, cat#: L1293, Sigma Aldrich), LMNB1 (1:1000, cat#: sc-365214, Santa Cruz), SUN1 (1:1000, cat#: NBP1-87396, Novus Biologicals), tubulin (1:1000, cat#: 3873S, Cell SignalingTechnology) Secondary antibodies: 680-or 790-conjugated Alexa Fluor (1:10000, cat#: A10043 and A11371, Thermo Fisher)

### NUP153 localization analysis

Cells were imaged with a 63X/1.40 objective with 0.2 μm z step. In Imaris, surfaces were created based on DAPI signal. All cutoff or missegmented nuclei were manually removed from analysis. Spots were created on NUP153 channel with estimated diameter of 0.2 μm and adjusted based on median intensity to minimize noise. Spots near surfaces were counted and divided by surface area.

### Lamina Gap Analysis

Cells were imaged with a 63X/1.40 objective with 0.2 μm z-step and processed with lightning deconvolution (Leica). An app was created in MATLAB (R2022b) for the quantification and characterization of lamina gaps. The algorithm that we developed to identify gaps in the 3D nuclear envelope relies on the comparison between the regular, cartesian distance transform map, and the geodesic distance transform map. The rationale is that the geodesic distance between a given pixel and a reference point is the same as the regular, cartesian distance only if the path is not constrained (i.e., a gap in the lamina). Finding equality between the two distance maps will thus reveal the gap areas, akin to rays of light coming out of a punctured sphere containing an illumination source. These rays of light are then projected onto the contour of the nucleus (defined by the perimeter of the DAPI signal) to extract the precise size and location of the gaps.

Briefly, Lamin and DAPI signals were segmented in 3D using standard procedures. After a slight dilation of the nucleus (DAPI) object, the perimeter pixels of the resulting object were defined. Erosion of the DAPI object was used as a reference/ seed for distance transform computations, and the complement of the binarized lamin signal was used to constrain the geodesic distance computation. The intersect between the dilated DAPI perimeter and the distance equality map defines the lamina gaps. Mis-segmented nuclei were removed from analysis based on nuclear volume and negative curvature of gaps. Gaps over 5 um^3^ were manually verified and resulted from merging multiple gaps nearby. These gaps were removed from analysis.

To quantify the local nuclear curvature associated with lamin gaps, we created a bounding volume (alphashape) that envelops the binarized DAPI signal with a cloud density of 10 points per μm^2^. Position, surface normals and curvature values were then extracted for each point using a 3-by-3 kernel. This analysis was implemented by the ‘findPointNormals()’ function^100^. Curvature values covering a given gap were then averaged.

### Statistics and quantification

Quantification of fixed image data generated by CellProfiler 4.2.4: Images corrected for background signal were analyzed for mean intensity RFP and GFP in the nucleus and cytoplasm using mask objects generated by rescaling intensity, down-sampling, median filtering, and adaptive Otsu thresholding of Hoechst (nucleus) and RFP (cytoplasm). Morphological measurements of the nucleus used for data filtering and comparisons were generated from the Hoechst mask. Rupture frequency for fixed images was determined either by data output from CellProfiler, in which rupture was determined if the nuc:cyto ratio of RFP exceeded one standard deviation above the median for the experimental control (RFP+), cyto:nuc ratio of GFP exceeded one standard deviation above the median for the experimental control (GFP+), or manually (as noted for validation) based on visual identification of nuclear RFP signal. Intensity outputs from CellProfiler were measured from eroded masks to reduce error from over-segmentation.

Quantification of rupture frequency and duration in time-lapse image analysis was done manually in ImageJ by visually identifying a rapid loss of compartmentalization and subsequent restoration of nuclear localization signal, as previously implemented^46^.

Hypergeometric enrichment analysis was completed in R 4.2.1 using the mysigdbr and clusterProfiler packages. For enrichment of statistically significant rupture frequency hits within the context of the screen, gene sets from the Broad Institute C5:GO database were first joined to the complete gene list for the siRNA screen, ensuring hit enrichment calculations were performed relative to the landscape of the screen. Enrichment scores were determined by dividing the gene ratio by the background ratio, and p-values determined by Fishers exact test as generated by the enricher function (enricher(gene = hits, TERM2GENE = screen)).

All statistical tests were conducted using R (version 4.2.1). Each dataset was completed with a minimum of N = 3 biological replicates. Pair-wise comparisons on categorical data were analyzed using Barnard’s exact test (“Barnard” package, R), comparing non-gaussian distributions utilized Wilcoxon rank sum test (base R), and gaussian distributions by Kolmogorov-Smirnov test (base R). Bonferroni correction for multiple testing applied as indicated for statistical analyses with greater than five comparisons. Tables for statistical analyses are included as tabs in the Supplementary Tables spreadsheet.

