## Supplementary material for "A high-content screen reveals new regulators of nuclear membrane stability": Table of Contents for Supplementary Movies and Tables

**Movie S1.** Shown are a rupturing (right) and non-rupturing (left) U2OS RuptR cells. Channel order = RFP-Cyto, GFP-Nuc, merge. Scale bar = 10 um. Time stamp = hh:mm. Movie speed = 7 fps.

**Movie S2.** False negative U2OS RuptR cell, with rupture occurring at 11 s. Although there is a slight increase in nucleus RFP, the mean intensity did not meet the threshold for this single frame rupture. Channel order = RFP-Cyto, GFP-Nuc, merge. Scale bar = 10 um. Time stamp = hh:mm. Movie speed = 1 fps.

**Movie S3.** False Positive U2OS RuptR cell. Channel order = RFP-Cyto, GFP-Nuc, merge. Scale bar = 10 um. Time stamp = hh:mm. Movie speed = 7 fps.

**Movie S4.** U2OS RuptR cell, with single frame small rupture occurring at 11 s. Channel order = RFP-Cyto, GFP-Nuc, merge. Scale bar = 10 um. Time stamp = hh:mm. Movie speed = 1 fps.

**Movie S5.** Shown is a mitotic U2OS RuptR cell, channel order RFP-Cyto, GFP-Nuc, merge. Scale bar = 10 um, time stamp = hh:mm, image speed = 7 fps.

**Movie S6.** Shown are nuclear ruptures in U2OS shLMNB1 2xRFP-NLS transfected with siCTRL (left) and siCTDNEP1 (right) siRNAs. Channel = RFP-NLS. Scale bar = 10 um, time stamp = hh:mm, image speed = 7 fps.

**Supplementary Tables**

Table S1. Statistics associated with Fig. 1F, live vs fixed cell analysis of RuptR

Table S2. Statistics associated with Fig. 2D, known siRNA pipeline validation

Table S3. Statistics associated with Fig. 3A, siRNA screen RFP+ hits analysis

Table S4. Statistics associated with Fig. 3B, RFP-Cyto hits validation

Table S5. Statistics associated with Fig. 3C, siRNA screen GFP+ hits analysis

Table S6. Statistics associated with Fig. 3D, GFP-Nuc hits validation

Table S7. Statistics associated with Fig. 3E, siRNA screen nucleus area analysis

Table S8. Statistics associated with Fig. 3F, siRNA screen nucleus solidity analysis

Table S9. Table of siRNAs included in screen

Table S10. Statistics associated with Fig. 4A, hypergeometric enrichment GOBP

Table S11. Statistics associated with Fig. 4B, hypergeometric enrichment GOCC

Table S12. Statistics associated with Fig. S4A, hypergeometric enrichment targeted vs random

Table S13. Statistics associated with Fig. S4B, hypergeometric enrichment screen bias

Table S14. Statistics associated with Fig. 5A, CTDNEP1 rupture frequency analysis

Table S15. Statistics associated with Fig. 5B, CTDNEP1 rupture duration analysis

Table S16. Statistics associated with Fig. 5C, CTDNEP1 micronucleus rupture analysis

Table S17. Statistics associated with Fig. 5E, S5D, S5E, CTDNEP1 nucleus morphology analysis

Table S18. Statistics associated with Fig. S5B, CTDNEP1-10 rupture frequency analysis

Table S19. Statistics associated with Fig. 6C, CTDNEP1 nucleus gaps proportions

Table S20. Statistics associated with Fig. 6D, 6E, CTDNEP1 nucleus gaps characteristics
