## Supplemental Figure 1 for "A high-content screen reveals new regulators of nuclear membrane stability"

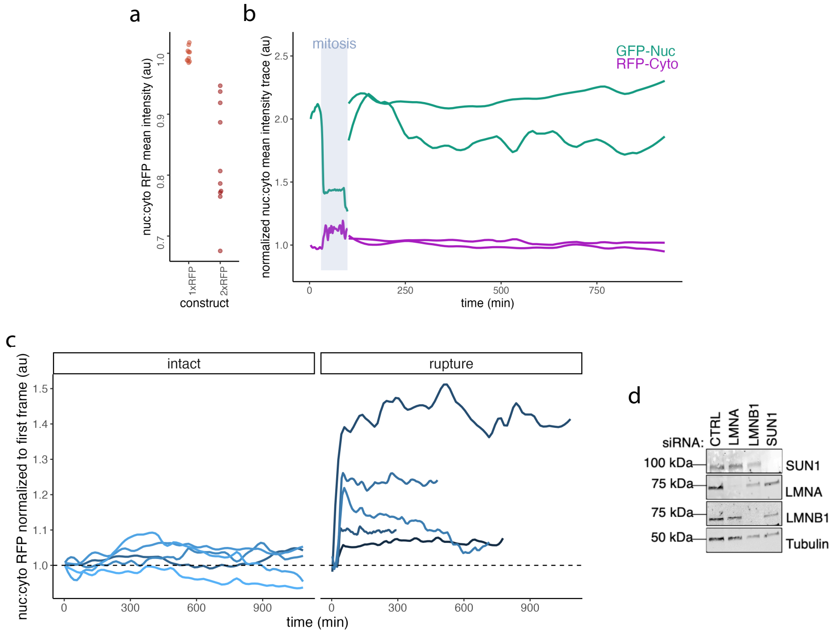


**Figure S1. a.** Comparison of nuc:cyto RFP mean intensity for 1x vs 2xRFP RFP-Cyto constructs after transfection in U2OS shLMNB1 GFP-Nuc cells. **b.** Nuc:cyto mean intensity over time for RFP-cyto and GFP-nuc resolution following mitosis. **c.** Traces of cells undergoing at least one rupture show sustained increase in nuclear RFP signal in rupturing but not intact cells **d.** Western blot indicating efficacy of validation siRNA knockdowns. Cells = U2OS RuptR.
