## Supplemental Figure 2 for "A high-content screen reveals new regulators of nuclear membrane stability"

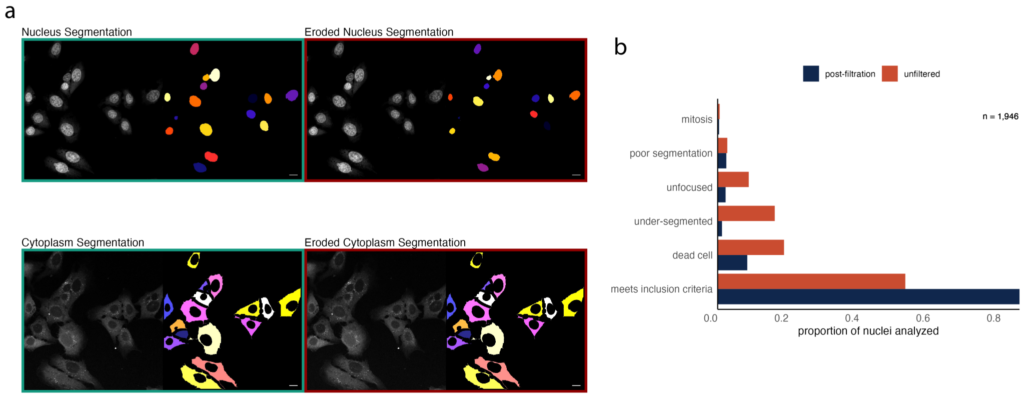


**Figure S2. a.** Representative images showing the segmentation for morphology output (left) and eroded for mean intensity measurements (right) **b.** Proportion of nuclei visually determined to be sufficient for analysis before and after applying morphology and intensity filters during the analysis pipeline.
