## Supplemental Figure 3 for "A high-content screen reveals new regulators of nuclear membrane stability"

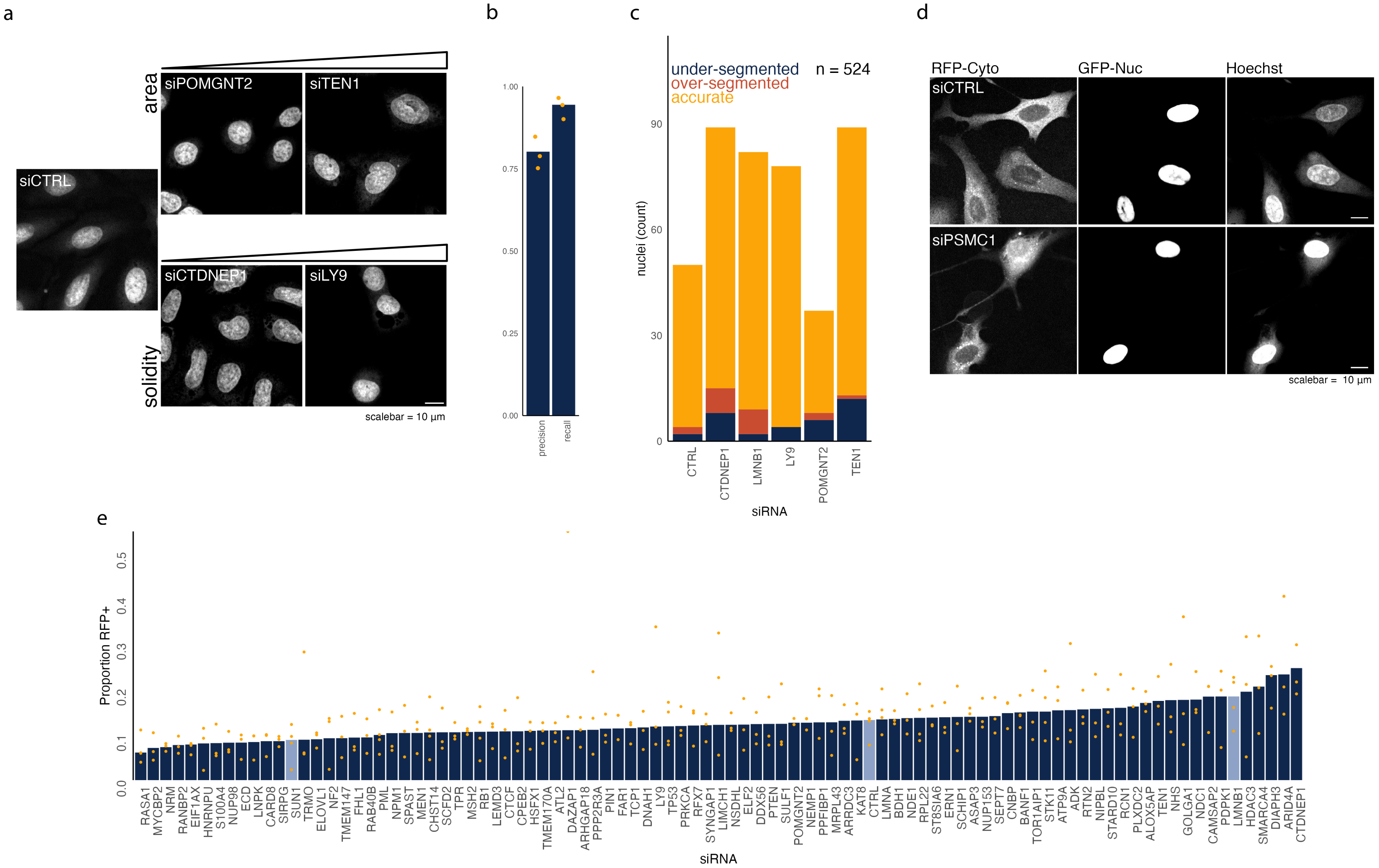


**Figure S3. a.** Representative images (Hoechst) of morphology screen hits. **b.** Manual precision/recall analysis for a subset of images from screen. **c.** Manual assessment of segmentation fidelity for indicated siRNAs **d.** Representative images of CTRL and siPSMC1 cells showing evidence of cell death, including cell rounding and increased Hoechst intensity. **e.** Full screen results showing the fold change nuc:cyto RFP ratio values over filter threshold (RFP+). Cells: U2OS RuptR.
