## Supplemental Figure 4 for "A high-content screen reveals new regulators of nuclear membrane stability"

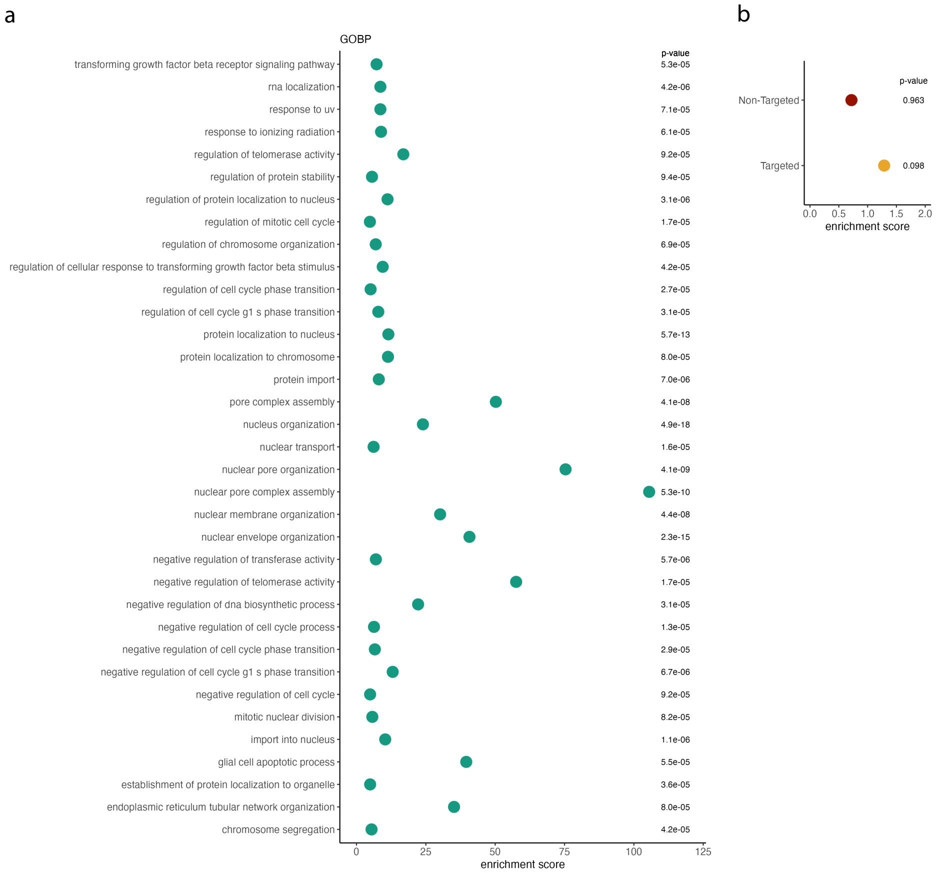


**Figure S4. a.** Enrichment of all siRNAs included in the screen relative to the GOBP gene set, p-value indicated on the right. **b.** Hypergeometric enrichment finds bias but not significant enrichment of siRNAs specifically selected for the screen in the screen hits. Stats: Table S12-13.
