## Supplemental Figure 5 for "A high-content screen reveals new regulators of nuclear membrane stability"

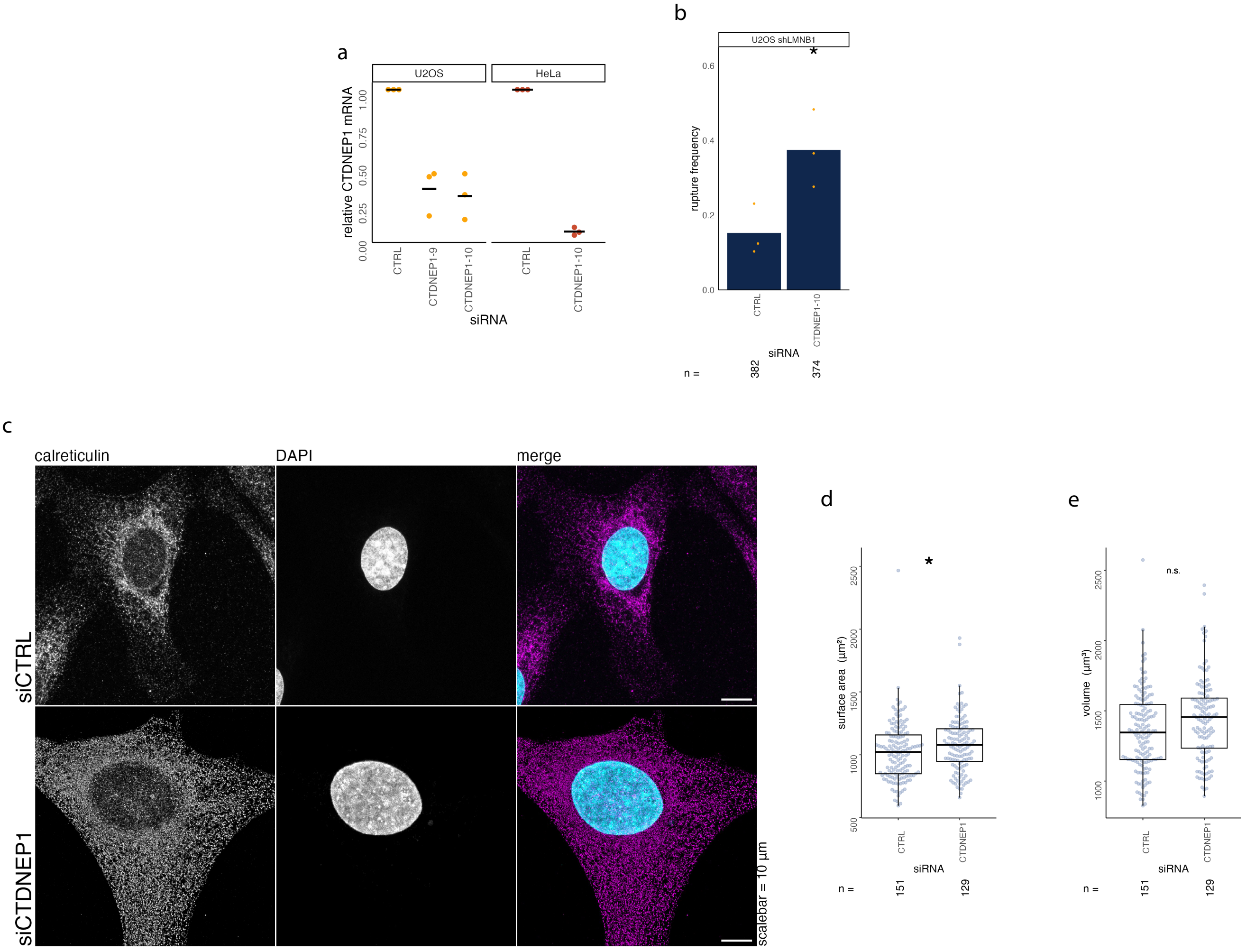


**Figure S5. a.** qRT-PCR results for CTDNEP1 siRNAs (9 and 10) in U2OS and HeLa cells **b.** Rupture frequency increases in U2OS shLMNB1 2xRFP-NLS cells following siRNA depletion of CTDNEP1 using a second siRNA (CTDNEP1-10) **c.** Representative image panel showing increased endomembrane system (calreticulin) in CTD-NEP1 depleted cells **d-e.** Distribution of nucleus surface area (d) and volume (e) in siRNA treated cells. * = $p < 0.05$, Cells: U2OS unless otherwise noted, Stats: Table S17-S18
